## Supplementary figures and images for "Population-level morphological analysis of paired CO_2_- and odor-sensing olfactory neurons in *D. melanogaster* via volume electron microscopy"

### Supplemental Figure 1

Figure 1_figure supplement 1

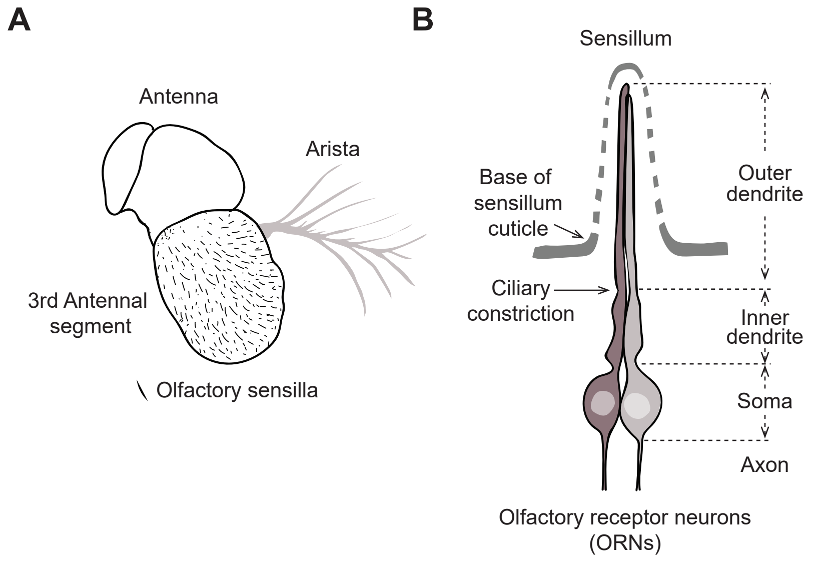

### Supplemental Figure 2

Figure 2_figure supplement 1

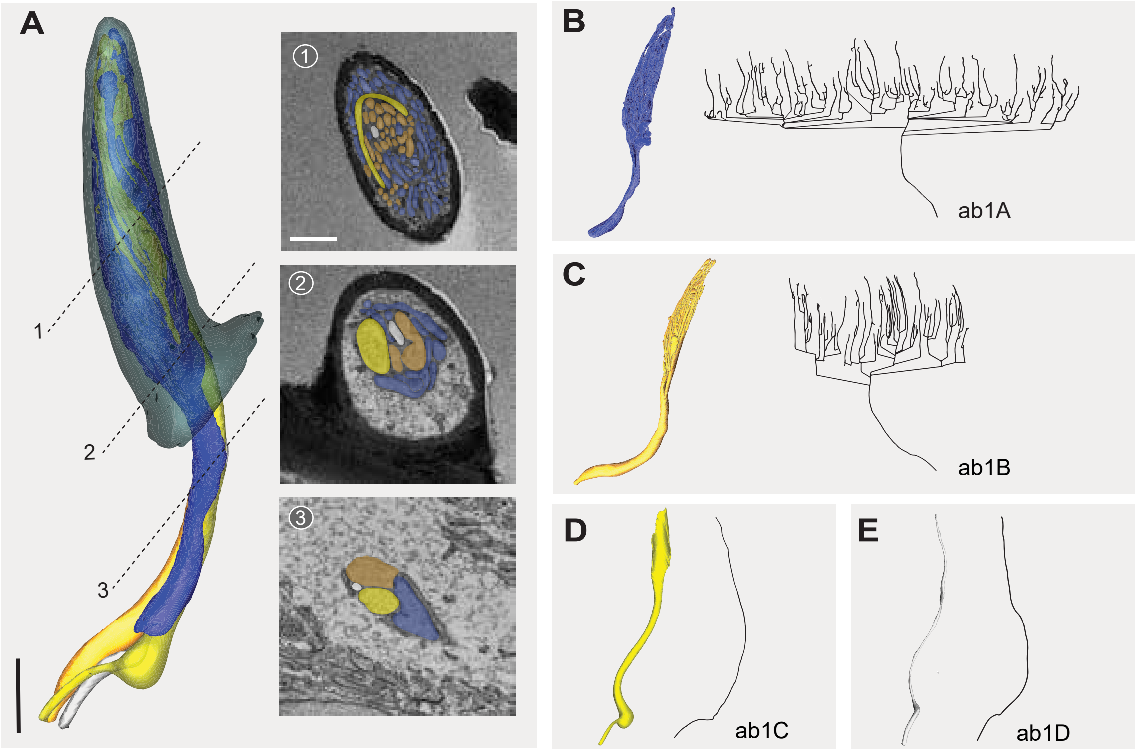
